# Protocol for Extracting Circulating Cell-Free DNA from Murine Saliva: Insights into Oral and Systemic Disease Research

**DOI:** 10.1101/2025.03.31.645839

**Authors:** Iram Elamin, Meghna S. Rao, Robert W. Figliozzi, Jessica Caroline Maahs, Matthew Balish, S. Victor Hsia, Ana Paula Piovezan Fugolin, Jiabing Fan

## Abstract

Circulating cell-free DNA (cfDNA) consists of small fragments of extracellular DNA from mammalian and bacterial cells found in bodily fluids such as blood and saliva, and it has been strongly recognized as a critical biomarker for various disease diagnoses, prognoses, and therapeutic monitoring. In this study, we present a reproducible protocol for efficiently isolating cfDNA from murine saliva using an innovative swabbing method in conjunction with the QIAamp MinElute ccfDNA Mini Kit. The quantification of isolated cfDNA is detected by a Qubit Fluorometer. Moreover, qualification assessment is conducted through BioAnalyzer analysis. This protocol facilitates research on saliva-derived cfDNA in the context of oral and systemic diseases in murine models.

**Graphical Abstract:** 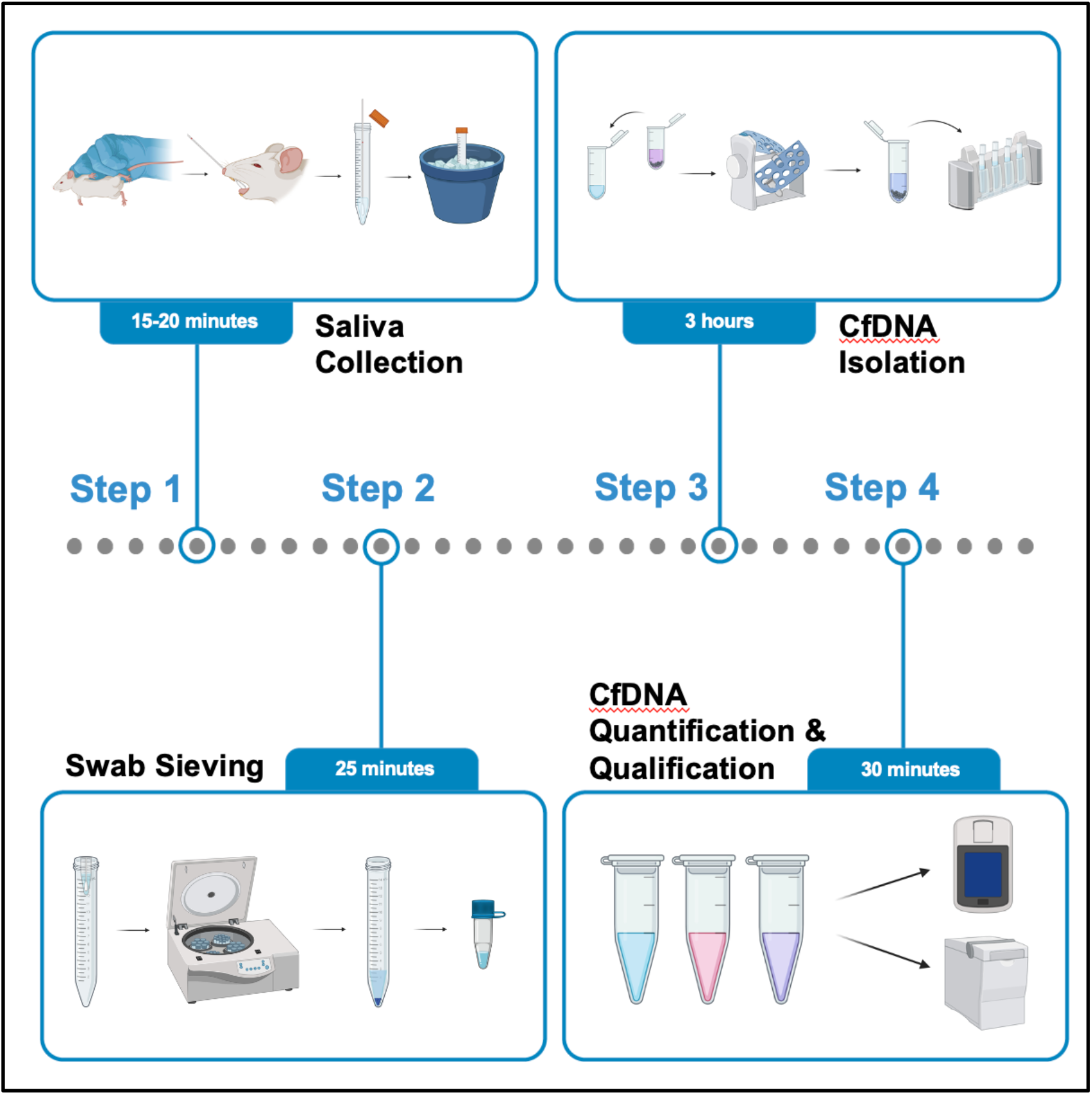

## BEFORE YOU BEGIN

Circulating cell-free DNA (cfDNA) are short fragments of extracellular DNA that circulate in various bodily fluids, including blood, saliva, urine, and cerebrospinal fluid (CSF).^1,2^ This cfDNA is released into the circulation through several cellular processes, most notably apoptosis, necrosis, and active secretion by living cells.^1,2^ Additionally, blood samples often contain circulating microbial cfDNA, which mainly originates from degraded microbial cells through processes such as NETosis, phagocytosis, and the Membrane Attack Complex.^3,4^ The presence of cfDNA in bodily fluids has attracted significant interest due to its potential as a non-invasive biomarker for disease diagnosis, monitoring treatment responses, and predicting disease progression.^5,6^ This is particularly relevant in the context of liquid biopsy technologies, which offer a minimally invasive approach to cancer detection and management, thus aligning with the principles of precision medicine.

Current research predominantly centers on the extraction and analysis of cfDNA from human biofluids such as plasma, serum, urine, CSF, and saliva.^7,8^ Each of these fluids has unique properties and clinical relevance. For example, cfDNA derived from plasma is often used in oncology to detect tumor-specific mutations, while urinary cfDNA has been studied for its potential in diagnosing urological cancers.^8-10^ Saliva, being an easily accessible biofluid, has emerged as a promising source for cfDNA analysis, particularly in the realm of oral and systemic diseases.^11,12^

In the field of salivaomics – an area dedicated to the study of salivary components, most investigations have focused on human samples and methodologies.^13^ However, research involving murine models provides invaluable insights that can improve studies related to human health, especially because it allows the induction of diseases and conditions in a controlled setting. When analyzing cfDNA from murine saliva, standard protocols often involve pharmacological stimulation to induce salivation. In various studies, pilocarpine, a muscarinic agonist, is widely used to significantly increase salivary secretion.^14-16^ These protocols generally yield approximately 70-200 µL of saliva over several minutes of stimulation, with samples collected from anesthetized mice.^14-16^ To ensure the welfare of the animals and the integrity of the samples, it is imperative to adhere to specific guidelines for saliva collection.^17,18^ Saliva sampling should be limited to volumes not exceeding 100 µL per collection to enhance the success of resuscitating the animals. Furthermore, to minimize stress and physiological impact, a minimum interval of one week should be maintained between successive saliva collections. Adhering to these guidelines is crucial, as deviations have been associated with a significant mortality rate of pilocarpine-induced mice.^18,19^

In this context, we present a protocol designed for the efficient isolation of cfDNA from a cohort of mouse saliva. This protocol aims to facilitate subsequent experimental investigations into the role of saliva-derived cfDNA in various biological and pathological processes, particularly those related to oral and systemic diseases.

### Institutional Permissions

All animal protocols were approved by the University of Maryland Eastern Shore Institutional Animal Care and Use Committee and were strictly in compliance with the Guidelines for the Care and Use of Laboratory Animals of the National Institutes of Health.

## KEY RESOURCES TABLE

| REAGENT OR RESOURCE | SOURCE | IDENTIFIER |
| --- | --- | --- |
| Mice Strain |  |  |
| C57BL/6J | The Jackson Laboratory | 000-664 |
| Chemicals or Reagents |  |  |
| QIAamp MinElute ccfDNA Mini Kit | Qiagen | 55204 |
| Qubit 1X dsDNA HS Assay Kit | Invitrogen | Q33231 |
| PBS (1X) | Corning | 21-040-CV |
| Tris-EDTA Buffer (pH 8) | VWR Life Sciences | 3247C241 |
| Absolute Ethanol | Fisher Scientific | A409-4 |
| Isopropanol | Honeywell | 34965-1L |
| Swabs | Fisher Scientific | 23-600-985 |
| Microcentrifuge tubes (1.5ml) | Fisher Scientific | 22431021 |
| Microcentrifuge tubes (0.5ml) | Fisher Scientific | 22431005 |
| Conical tubes (15ml) | Fisher Scientific | 12-565-269 |
| Beaker | Fisher Scientific | FB100250 |
| Aluminum Foil | Fisher Scientific | 15-078-292 |
| Agilent High Sensitivity DNA Kit | Agilent Technologies | 5067-4626 |
| Other |  |  |
| Centrifuge | ThermoFisher Scientific | 75004261 |
| Rotor | ThermoFisher Scientific | 75003629 |
| Rotor | ThermoFisher Scientific | 75003652 |
| Centrifuge | ThermoFisher Scientific | 75002461 |
| Minicentrifuge | Fisher Scientific | S67601B |
| Analytical Balance | Sartorius | BCE64-1S |
| Tube Rotator | Fisher Scientific | 88861051 |
| Fluorometer | Invitrogen | Q33226 |
| Thermal Mixer | ThermoFisher Scientific | 13687711 |
| Magnetic Rack | Qiagen | 399911 |
| Incubator | New Brunswick Scientific | INNOVA 4335 |
| Vortex | Fisher Scientific | 14-955-152 |
| BioAnalyzer 2100 | Agilent Technologies | N/A |

## STEP-BY-STEP METHOD DETAILS

### Saliva Collection

**Timing:** 15-20 minutes

1. Pipette 1 mL of refrigerated phosphate-buffered saline (PBS) into a 15 mL conical tube.
2. Gently restrain the mice using a standard rodent handling technique.
3. Insert a nasopharyngeal swab into the oral cavity of each mouse and gently rotate it for 60 seconds, ensuring contact under the tongue, in each cheek cavity, and along the front edges of the oral cavity.
4. Place the nasopharyngeal swab into the 15 mL conical tube and repeat step 3 to collect sufficient saliva samples (**Figure 1**).

**Figure 1:**
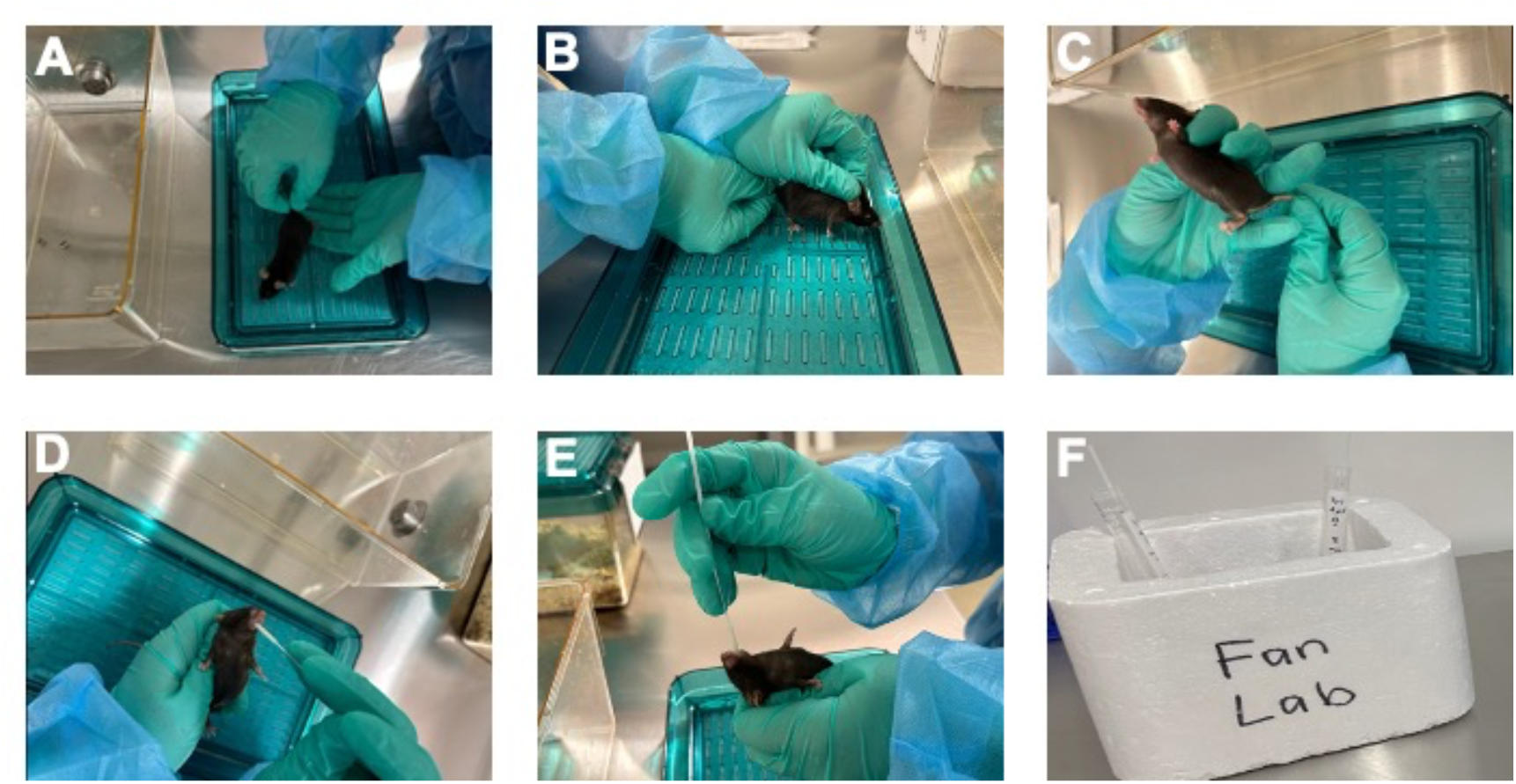
(A-F) The schematic diagram of collecting saliva from mouse (steps 1-4).

**Note:** Prior to saliva collection, PBS should be stored at 4°C. All tubes must be kept on ice during swabbing and stored at 4°C; do not store for more than one day. Ensure you wear appropriate protective gear when handling animals. Care should be taken not to insert the swab beyond the tongue or into the throat or esophagus. Do not allow the mouse to chew off and swallow parts of the swab. Extreme caution must be exercised to avoid restricting the respiration of mouse or pinching any muscle, bone, vital tissue, or nerve. At any sign of distress, the animal should be gently returned to its housing, and a new animal should be selected for the procedure.

### Swab Sieving

**Timing:** 25 minutes

5. Pierce the bottom of a 1.5 mL DNA LoBind PCR clean tube using a 16-gauge needle.
6. Place the swabs (soaked in PBS prior to use) into the 1.5 mL tube and trim off the excess (**Figure 2**).
7. Return this assembly to the 15 mL conical tube.
8. Centrifuge the tubes for 10 minutes at 1900g (4000 rpm) at 4°C.
9. Carefully collect the supernatant (sample) without disturbing the pellet. Transfer the supernatant into a fresh 1.5 mL tube placed on ice and discard the pellet (debris).
10. Centrifuge collected supernatant at 10,000g (9500 rpm) for an additional 10 minutes at 4°C to remove other residual cell debris.
11. Transfer the supernatant into freshly labeled 1.5 mL tubes and keep the tubes on ice until further processing.

**Figure 2:**
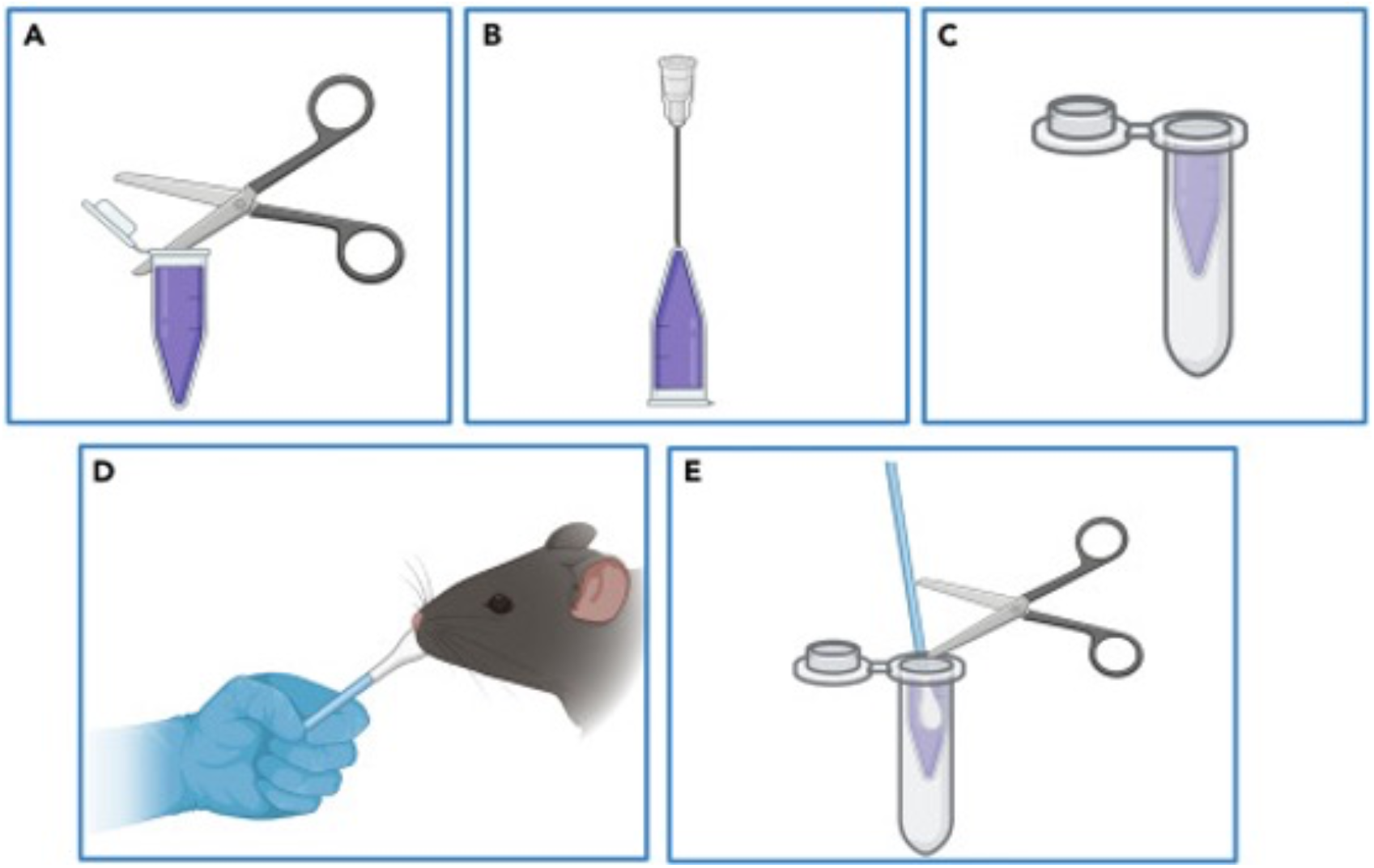
(A-E) The schematic diagram of the placement of swabs into 1.5 mL tubes for the initial centrifugation step in swab sieving (steps 5 and 6).

**Note:** Keep the tubes open during centrifugation, as closing them may create a vacuum. Weigh an empty tube beforehand to determine its standard weight. After the first centrifugation, the supernatant is collected in a fresh tube and recorded in volume and weight. Calculate the sample weight by subtracting the standard tube weight from the total weight of the tube containing the sample.

### Cell-Free DNA Isolation Using the QIAamp MinElute ccfDNA Mini Kit

**Timing:** 3 hours

12. Prepare a mixture of 55 µL proteinase K, 150 µL bead binding buffer, and 30 µL magnetic beads for every 1 mL of sample.
13. Add 235 µL of the mixture to each sample tube.
14. Wrap the tubes in aluminum foil and place them in an end-over-end centrifuge for 10 minutes at room temperature (configure the instrument to the default rotation setting).
15. Centrifuge the tubes briefly for 10-20 seconds.
16. Place the tubes in a magnetic rack for 1 minute, ensuring that the metal beads are aggregated.
17. Discard the supernatant (clear solution) by carefully positioning the pipette tip against the opposite wall of the tube, away from the beads, and collect the supernatant (keep the cap open during this step).
18. Add 200 µL of the bead elution buffer to the tubes containing the bead pellets.
19. Mix and rinse the bead pellets from the tube wall, then resuspend them in the bead elution buffer.
20. Vortex the mixture thoroughly. Perform a short centrifugation to gather the resuspension, gently remixing the pellet without splashing the solution on the tube walls.
21. Transfer the suspension into a 2 mL bead elution tube.
22. Wrap the tubes in aluminum foil and place them on a shaker at 300 rpm for 5 minutes at room temperature.
23. Place the tubes on the magnetic rack until the solution is clear.
24. Transfer the supernatant into a fresh bead elution tube and add 300 µL of the ACB buffer to the supernatant. Vortex the mixture.
25. Transfer the mixture into a MinElute column and centrifuge for 1 minute at 6000g (7900 rpm) at room temperature.
26. Place the column in a new 2 mL collection tube and discard the bottom tube containing the flow-through.
27. Add 500 µL of ACW2 buffer to the column.
28. Centrifuge for 1 minute at 6000g (7900 rpm) at room temperature.
29. Discard the flow-through and place the column in a new 2 mL collection tube.
30. Centrifuge the column for 3 minutes at 20,000g (14,400 rpm) at room temperature.
31. Place the column in a 1.5 mL collection tube.
32. Incubate this assembly in an oven for 3 minutes at 56°C (keep the cap open to dry the membrane).
33. Add 25 µL of ultra-clean water to the center of the column for re-elution, close the lid, and incubate at room temperature for 1 minute.
34. Centrifuge for 1 minute at 20,000g (14,400 rpm) to elute the nucleic acids.

**Note**: Prepare a stock solution for multiple samples, including an additional excess to minimize pipetting errors. For example, to prepare a stock solution for 10 samples, combine 605 µL of proteinase K, 1650 µL of bead binding buffer, and 330 µL of magnetic beads (step 12). Repeat step 34 after transferring the solution from step 33 to the column for maximum elution. After this step, store the tubes on ice or at −80°C for long-term preservation. Limit the sample input volume to 1-4 mL.

### Cell-Free DNA Quantification & Qualification Assessment

a. **Cell-Free DNA Quantification Using Qubit 1xdsDNA HS Assay Kit**

**Timing:** 5 mins

35. Add 198 µL of working reagent to a 0.5 mL centrifuge tube, then introduce 2 µL of the cfDNA sample.
36. In a new 0.5 mL centrifuge tube, add 190 µL of working reagent followed by 10 µL of the negative standard.
37. In another new 0.5 mL centrifuge tube, add 190 µL of working reagent followed by 10 µL of the positive standard.
38. Incubate the mixtures for 2 minutes at room temperature.
39. Read the samples using a Qubit Fluorometer.

**Note:** Turn off the light in the LAF hood before starting this experiment, as the working reagent is light-sensitive. Vortex all mixtures thoroughly before incubation. Make sure to select the correct settings (dsDNA > 1x dsDNA High Sensitivity) and sample volume (2 µL). Begin by analyzing standards, followed by the samples. The concentration of cfDNA from our approach is approximately 1000 ng/mL.

b. **Cell-Free DNA Qualification Assessment**

**Timing:** 25 mins

40. The isolated cfDNA is assessed for quality using the Agilent BioAnalyzer 2100 with the Agilent High Sensitivity DNA Kit. Load the prepared samples onto the BioAnalyzer and run the assay under standard conditions for cfDNA analysis.
41. The cfDNA size distribution is further analyzed, focusing on fragments within the 50–300 bp range to determine the proportion of small DNA fragments.

**Note:** Follow the manufacturer’s protocol for sample preparation, including proper dilution if necessary. The expected peak for cfDNA should be around 150 bp, corresponding to mononucleosomal DNA. Larger fragments (>300 bp) may indicate contamination with high-molecular-weight genomic DNA.

### Expected Outcomes

cfDNA has emerged as an important biomarker for disease diagnostics, prognostics, and therapeutic monitoring. Analyzing cfDNA enhances our understanding of disease mechanisms and provides a foundation for developing effective diagnostic tools and treatments. A protocol has been established for the efficient isolation of cfDNA from mouse saliva. The innovative swabbing method for saliva collection from live mice significantly enhances the convenience and effectiveness of cfDNA extraction (**Figure 1 and 2**). This approach eliminates the need for drug-induced salivation, reducing stress and discomfort for the animals while fostering a more natural and less invasive experimental environment. This methodology enables a thorough investigation of saliva-derived cfDNA in relation to oral and systemic diseases, offering new insights into disease pathology and potential therapeutic strategies.

### Limitations

While the established protocol efficiently isolates cfDNA from mouse saliva, the quality and quantity of the isolated cfDNA are influenced by several factors. First, a second centrifugation step is necessary to eliminate residual debris from saliva; failure to perform this step may contaminate the isolated cfDNA with proteins and other contaminants (**Figure 3**). Second, using EDTA in saliva collection can reduce cfDNA degradation but may compromise its purity. In contrast, using PBS may enhance cfDNA purity but increases the risk of degradation (**Figure 4**). Third, mouse saliva collected in PBS should not be stored at 4°C for more than 48 hours. This limitation may increase the workload, as cfDNA isolation from mouse saliva must be performed promptly.

**Figure 3:**
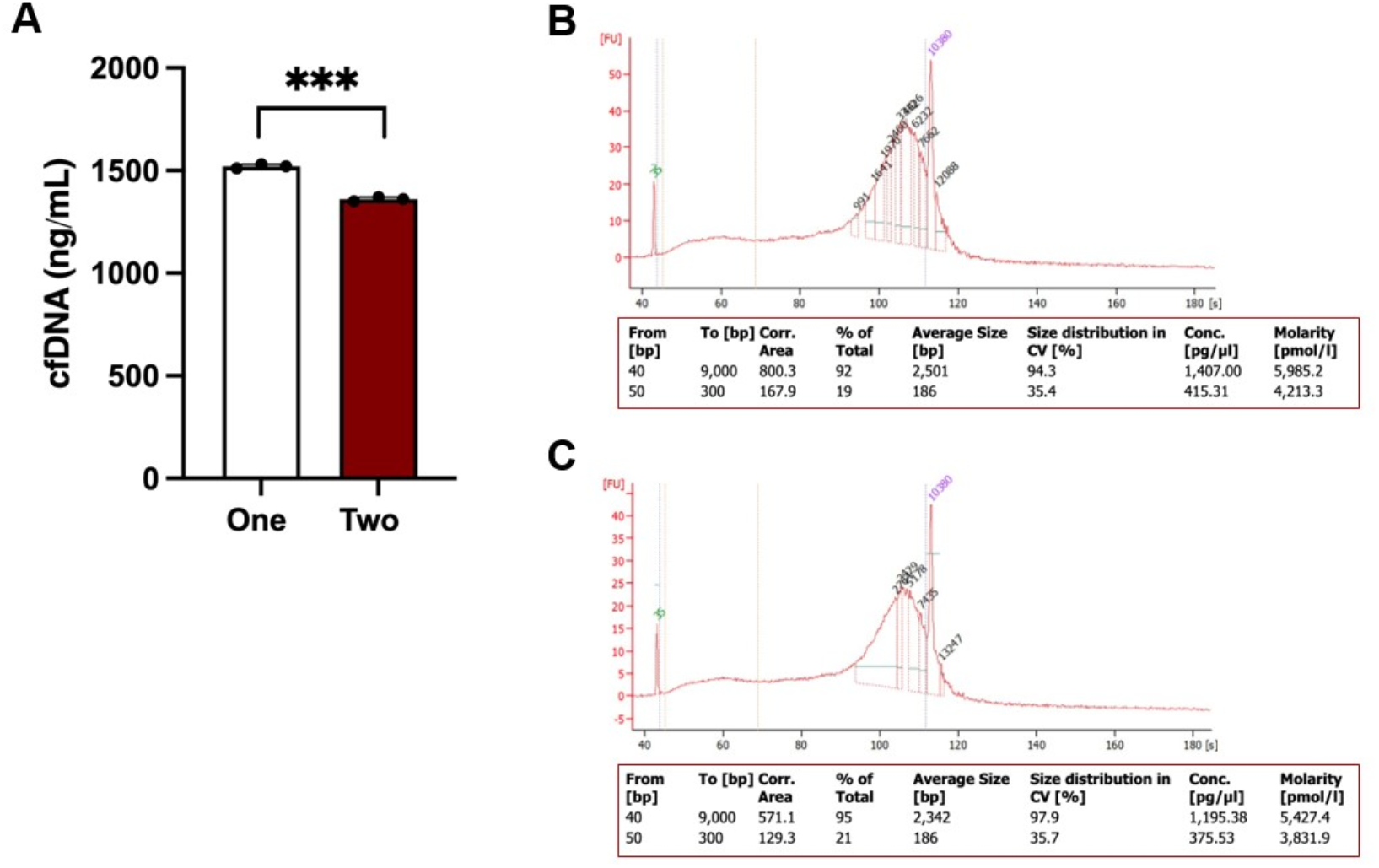
cfDNA isolation from saliva using EDTA with one- and two-time centrifugation. (A)Yield of cfDNA collected from saliva via one- and two-time centrifugation, with statistical significance indicated (***p < 0.001). (B, C) Quality assessment of cfDNA collected through one (B)and two (C) centrifugation steps was performed using the BioAnalyzer 2100 with the Agilent High Sensitivity DNA Kit. A size range of 50-300 bp was established to quantify the percentage of small DNA fragments.

**Figure 4:**
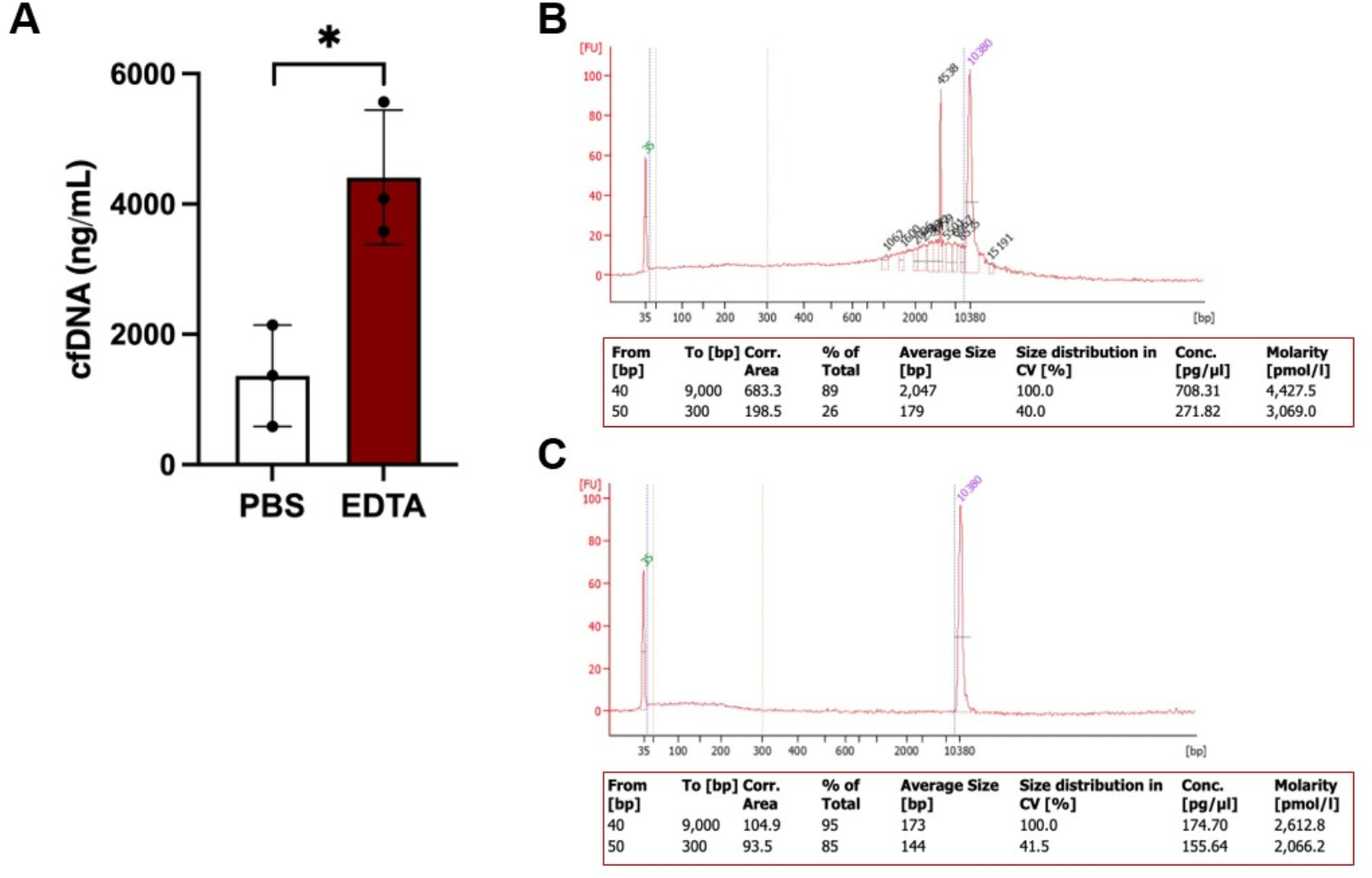
Quality assessment of cfDNA isolated from saliva using EDTA and PBS. (A) Yield of cfDNA collected from saliva with EDTA and PBS, with statistical significance indicated (*p < 0.05). (B, C) Quality assessment of cfDNA from saliva using EDTA (B) and PBS (C) was performed using the BioAnalyzer 2100 with the Agilent High Sensitivity DNA Kit. A size range of 50-300 bp was established to quantify the percentage of small DNA fragments.

Additionally, the volume of saliva that can be collected from mice is limited for each swab sample. Approximately 10-15 swab samples should be pooled for a single isolation, which restricts the overall yield of cfDNA due to the small total volume. Furthermore, frequent saliva collection can be time-consuming and may cause discomfort or potential oral damage to the mice. A minimum interval of approximately half an hour is required between collections to ensure sufficient saliva production. The choice of a swab is also critical. Cotton swabs readily adhere to saliva but can be challenging to transfer to collection tubes. In contrast, nylon flocked swabs are more efficient for transferring the collected saliva.

### Troubleshooting

**Problem 1**

### Buccal Swabbing

#### Potential Solutions

To perform buccal swabbing, securely hold the animal in your dominant hand. Use your non-dominant hand to select a sterile collection device—either a cotton swab for initial experiments or a nylon flocked swab for the optimized protocol. Carefully insert the swab into the slightly open mouth of the animal, ensuring that the tip does not touch the back of the throat or the esophagus. Rotate the swab to ensure it contacts all surfaces inside the mouth, including the teeth and side pockets, extending up to the nasal area. Limit the total contact time to 60 seconds, after which the animal should be promptly returned to its housing.

**Problem 2**

### Low saliva yield observed when using cotton swabs

#### Potential Solutions

Consider using nylon flocked swabs for saliva collection to increase yield.

**Problem 3**

### The PBS solution failed to preserve cfDNA in the saliva samples (related to steps 1-4)

#### Potential Solution

To minimize cfDNA degradation, cool the PBS solution before use and process the samples within 48 hours. Additionally, ensure that the samples are consistently kept on ice rather than at room temperature.

**Problem 4**

### Magnetic beads settle at the bottom of the tube (related to step 12)

#### Potential Solution

Vortex the magnetic beads thoroughly and use them immediately after mixing. Avoid leaving the beads undisturbed for more than two minutes.

**Problem 5**

### Disruption of the magnetic beads while discarding the supernatant (related to step 17)

#### Potential Solution

Keep the tubes on the magnetic rack. When discarding the supernatant, gently position the pipette tip against the opposite side of the tube, away from the beads, to prevent disturbance.

**Problem 6**

### Beads disintegrate when collecting the supernatant (related to step 24)

#### Potential Solution

Ensure the tube remains in contact with the magnetic rack and confirm bead formation by observing the clear solution. Carefully position the pipette tip on the opposite side of the tube, away from the beads, and transfer the supernatant to a fresh tube.

## RESOURCE AVAILABILITY

### Lead contact

Further information and requests for resources and reagents should be directed to and will be fulfilled by the lead contact, Jiabing Fan.

### Materials availability

This study did not generate new unique reagents.

### Data and code availability

- This paper does not report original code.
- Any additional information required to reanalyze the data reported in this paper is available from the lead contact upon request.

## ACKNOWLEDGMENTS

This work was supported by grants from the National Institutes of Health (R03 DE030539 to J.F.), and Osteo Science Foundation (to J.F.).

## AUTHOR CONTRIBUTIONS

Conceptualization, A.P.P.F and J. F.; methodology, I.E., M. S.R., R. W. F., A.P.P.F and J. F.; investigation, I.E., M. S. R., R. W. F., J. C. M., M. B and S. V. H. writing – original draft, I.E., M. S. R., R. W. F., A.P.P.F and J. F.; writing – review and editing, A.P.P.F and J. F.; funding acquisition, J.F.; supervision, A.P.P.F. and J.F.

